# Affordable Remote Monitoring of Plant Growth and Facilities using Raspberry Pi Computers

**DOI:** 10.1101/586776

**Authors:** Brandin Grindstaff, Makenzie E. Mabry, Paul D. Blischak, Micheal Quinn, J. Chris Pires

## Abstract

- *Premise of the study:* Environmentally controlled facilities, such as growth chambers, are essential tools for experimental research. Automated remote monitoring of such facilities with low-cost hardware can greatly improve both the reproducibility and the accurate maintenance of their conditions.
- *Methods and Results:* Using a Raspberry Pi computer, open-source software, environmental sensors, and a camera, we developed a cost-effective system for monitoring growth chamber conditions, which we have called ‘GMpi.’ Coupled with our software, *GMpi_Pack*, our setup automates sensor readings, photography, alerts when conditions fall out of range, and data transfer to cloud storage services.
- *Conclusions:* The GMpi offers low-cost access to environmental data logging, improving reproducibility of experiments, as well as reinforcing the stability of controlled environmental facilities. The device is also flexible and scalable, allowing customization and expansion to include other features such as machine vision.

## INTRODUCTION

Growth chambers play an important role in plant science and agronomics. It is vital that these chambers provide and maintain constant growing conditions such as light, temperature, and humidity in order to reduce variables that could bias experimental data. However, this is not always easy to do. The environmental parameters of growth chambers can fluctuate, and this can impede reproducibility in future experiments. In order to compensate for this issue, researchers typically will randomize placement of plants in the growth chamber and perform replications of the experiment. To make certain that the environmental parameters inside a growth chamber are within the required parameters, the best solution is to monitor and record them. This information can then be used to increase repeatability for future experiments and provide researchers with real time information about the conditions their plants are experiencing.

Single board computers paired with open source software provide the opportunity to develop such a system of growth chamber monitoring. The variety of different sensors and single board computers afford a high degree of flexibility and can be used in many different applications, such as IoT (internet of things) and otherwise. IoT devices are internet-connected “things” that are capable of collecting data and sending it through the internet. They can also be integrated to build scalable networks of reconfigurable computers capable of environmental monitoring (Ferdoush & Li 2014) or used for other tasks that can be invaluable in our data-driven field. Monitoring systems developed using this technology are most prevalent in agriculture. Two of the most popular applications are selective irrigation and predictive analytics, both of which can improve productivity and the efficiency of water management.

One example of such a system was developed by Shah & Bhatt (2017). Their system offers a means to monitor the conditions of a greenhouse using a Raspberry Pi, a temperature/humidity sensor, and a soil moisture probe that can autonomously log data to a cloud server over the internet. It can also be used to automate irrigation. Another recent application using Raspberry Pi’s is the monitoring of growing conditions for lettuce (*Lactuca sativa*). Here researchers used a network of Raspberry Pi computers equipped with sensors for light, temperature, and humidity as well as a real-time clock and external battery to monitor the plants (Cabaccan et al. 2017). These two applications provide excellent examples of the scalability of this technology as well as its ability to run on an independent power source for a period of time, giving it the flexibility needed to be used in places with limited access to electricity.

In the laboratory setting, Raspberry Pi’s are being used as a cost effective way improve the fidelity of controlled environments, such as incubators and growth chambers, that would have previously been too costly to monitor. In some cases, these monitors can detect system failures that would otherwise have gone unnoticed. Such was the case with Gurdita et al. (2016) when they developed a Raspberry Pi-based device to monitor an incubator and found an environmental control failure that the monitoring equipment in the incubator failed to detect. They then used the collected data to obtain a warranty replacement of the incubator’s control electronics. Not only does this technology enable new means and methods of data collection, it also offers a cost-effective way to improve existing monitoring systems through redundancy or improved precision depending on the system.

Using the Raspberry Pi as a platform, we have developed the Growth Monitor pi (GMpi; **Figure 1)** to be an affordable way to monitor plants in growth chambers and greenhouses. The small, credit card-sized computer, paired with a mobile phone-sized camera and various inexpensive sensors with access to the internet, allows for precise monitoring of experiments at low cost. This low cost point is also due in no small part to the free open-source softwares it is dependent on. This includes our software package, *GMpi_Pack*, which was written to automate setup and data collection for the GMpi. In addition to monitoring the plants, this device is capable of functioning autonomously, alerting users when an environmental parameter is out of a predetermined range, as well as building a large dataset of the conditions within the growth chamber over a user-designated period of time. The device we have developed stands out from other devices described above because our *GMpi_Pack* software combines multiple features, such as sensing (temperature, humidity, and light intensity), cloud storage, image capture, and alerts onto a single platform. It relies heavily on software already available from the open source community and is meant to illustrate how this technology can be developed and used by researchers who may not be as familiar with software engineering. With the detailed protocol below, we hope that this type of monitoring and sensing can now be made accessible to anyone.

**Figure 1:**
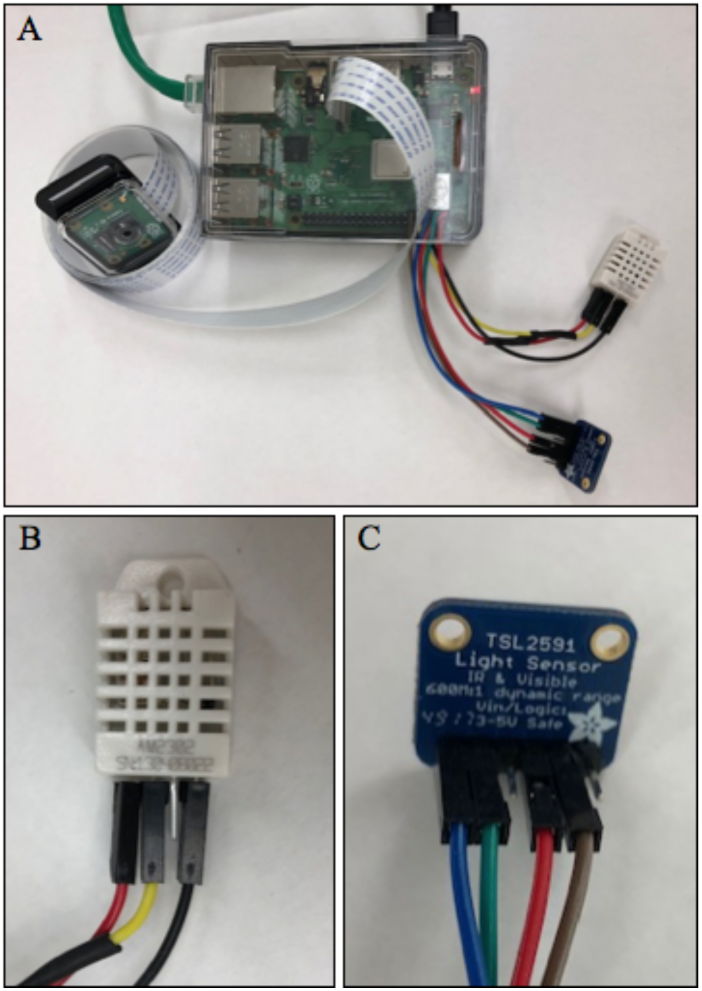
Images of the GMpi. (A) Photograph of the complete GMpi setup with all peripheries attached. (B and C) Close up photos of the temperature and humidity sensor (DHT22) and light intensity sensor (TSL2591).

## METHODS AND RESULTS

### Raspberry Pi and Peripheries

The growth chamber monitoring system was developed using a Raspberry Pi Model B+ single board computer installed with the Raspbian Stretch 4.14.50-v7+ operating system, but the most current version is recommended. We chose to use the Raspberry Pi over similar devices because of its low cost and because it is a commonly used computer with a host of software, information, and tutorials from the open source community. The model B+ was chosen over other models of the Raspberry Pi because it has 1 gigabyte of RAM and a processor rated at 1.4GHz. These resources are important as they afford performance overhead for prototyping, however, future models may be built with less powerful single board computers depending on one’s performance needs. A temperature and humidity sensor (DHT22; AdaFruit Industries LLC) was used to collect data about the humidity and temperature inside the growth chamber. The DHT22 model was chosen over the DHT11 due to its capability of detecting a higher range for both temperature and humidity readings and offering more accuracy in temperature readings (±0.5 °C compared to ±2 °C). DHT sensors are also useful as they are equipped with a capacitive humidity sensor and a thermal resistor. To collect data about light intensity in both full spectrum and infrared wavelengths, the TSL2591 (AdaFruit Industries LLC) sensor was used. This sensor is capable of detecting light between the ranges of 0.000118 to 88,000 Lux (lumens per square meter), allowing us to detect minute changes in light intensity in both the visible and infrared spectra. Although this light sensor is able to detect subtle changes in light intensity, the data collected is a calculated value (Lux), rather than a measured value, and the calculation is done using a function within the Python library for this device. It is effective for the purpose of detecting a change in light intensity, however, its measurements should not be considered a precise measurement. For that purpose a different sensor with a well documented calibration protocol is recommended. Lastly, a camera was connected to relay visual information to the end user over the internet, allowing for remote observation of plant growth.

### Connecting Sensors

To connect the different sensors to the Raspberry Pi, expertise in soldering is required. We enlisted the help from our department’s shop to achieve this. To connect the temperature and humidity sensor (DHT22), the first pen from the right was connected to a 3.3V general purpose input/output (GPIO) pin. The second pin was then connected to a data pin, the third was not used, and the fourth was connected to a ground (**Figure 2**). A 10KO resistor was soldered between the power wire and data wire to allow data output (**Figure 2**). For the light sensor (TSL2591), the first pin labeled “Vin” was connected to a 3.3V GPIO pin, then the pin labeled “GND” was connected to a ground pin (**Figure 2**). This device uses a I2C serial protocol in order to communicate with devices so the pins labeled “SDA” and “SCL” were connected to pins 3 and 5 of the Raspberry Pi. Finally, the camera was connected to the CSI-2 port behind the ethernet port via a ribbon cable. The ribbon cable provides additional flexibility in the placement of the camera for optimal viewing of plants.

**Figure 2:**
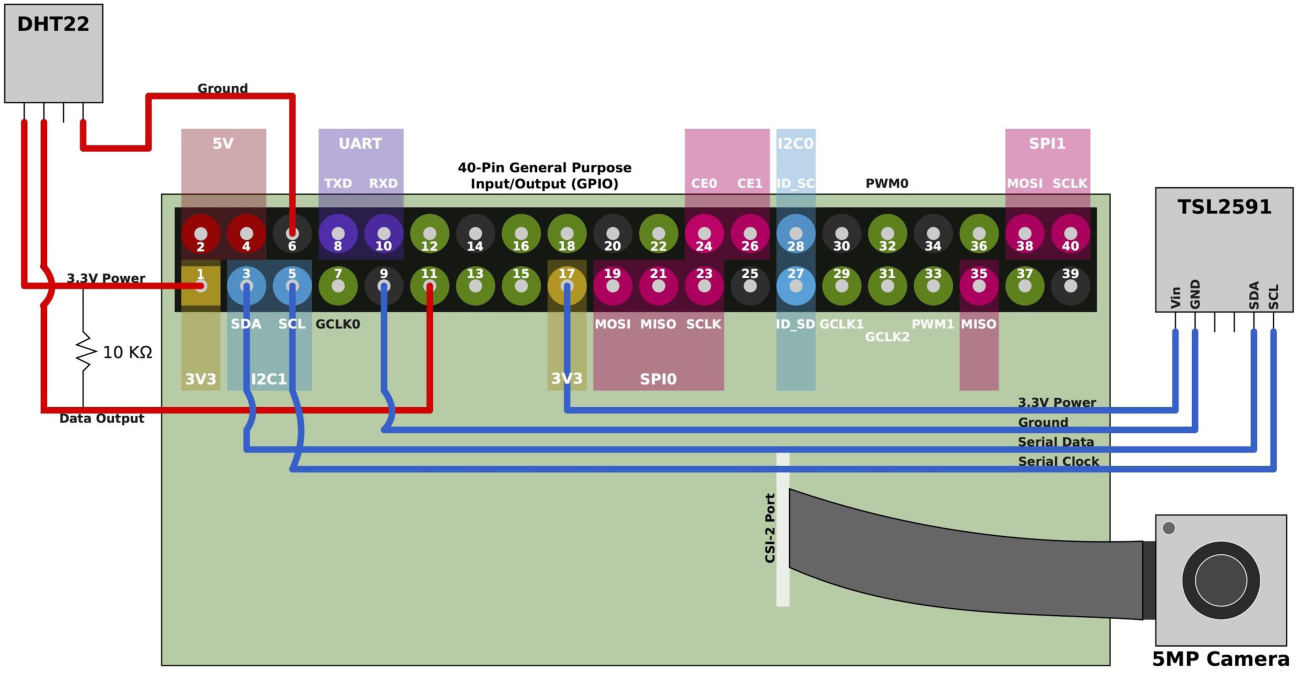
A visual diagram of the connections between the Raspberry Pi and the attached peripheries (above) and the Raspberry Pi GPIO pinout (below). Illustrated are the connections for the light intensity sensor (TSL2591) to GPIO the pins; pin 3 for SDA (serial data), pin 5 for SCL (serial clock), pin 9 for ground, and pin 17 for 3.3v power. It also illustrates the connection of the temperature and humidity sensor (DHT22) to the GPIO pins: pin 1 supplying 3.3v power, pin 6 for ground, and pin 11 for data. Importantly, it shows where the resistor should be soldered between the power and data wire for the temperature and humidity sensor (DHT22). Finally, it shows where the ribbon cable is from the camera board is connected to the Raspberry Pi.

### Uploading to a Cloud Drive Service

In order to upload the resulting sensor data to a cloud service, we used a third party, open source application called Rclone (github.com/pageauc/rclone4pi). This application can be easily configured to connect to several popular cloud storage services such as Google Drive, Box, Dropbox, etc. We elected to use Google Drive because it is a free service and it is a popular option that many people use. After you have adjusted your rclone configuration settings to your liking, you are given a link that allows a connection for this software to your cloud service. Uploading the collected data and images is handled by the *UploadFile()* and *UploadFile2()* functions in *GMPi_Pack.py* library (discussed below). Both of these scripts are called in *packageTest.py, picSnap.py*, and *upload.py* files and can be run automatically using the Unix tool cron.

### Remote Monitoring

A unique addition to the GMpi is its ability to notify users if a growth chamber or greenhouse falls out of a specified range for temperature, humidity, or light. We take advantage of the popular platform Slack (https://slack.com/), using the incoming webhooks function to set up remote notifications (**Figure 3**). This allows notifications to be sent to a configured Slack channel when thresholds are hit. As the use of Slack to communicate with research groups increases, it makes sense to have equipement be able to communicate via Slack as well. Using Slack Incoming Webhooks allows for end-users to configure what messages will generate a notification being sent from Slack to their device(s) that access Slack. It is also possible to ‘mention’ users directly in the Slack webhook message so that specific users get a notification (slack notification settings permitting) or one can configure a channel mention and it will notify all users in the channel (again, notification settings permitting).

**Figure 3:**
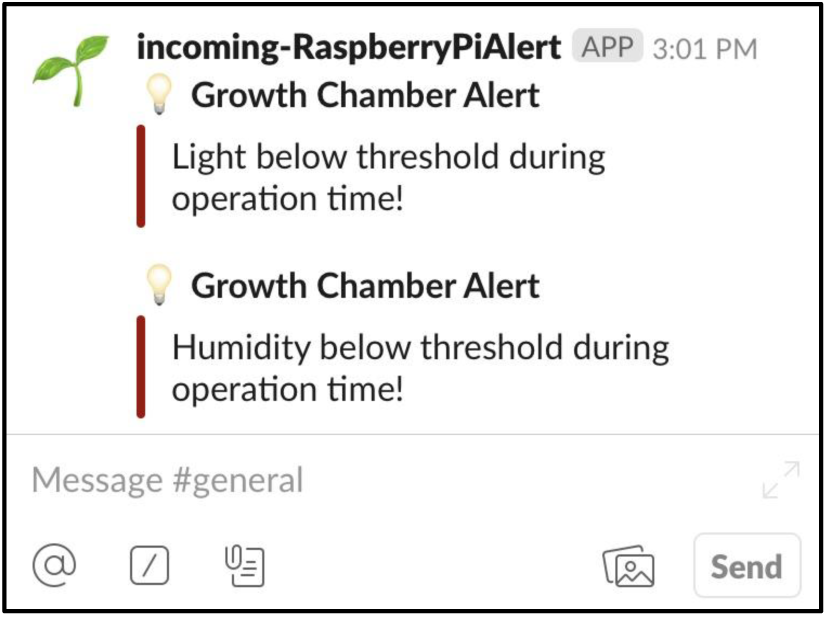
A snapshot of what an incoming alert from the GMpi looks like on Slack. This will alert all members of this workspace that the growth chamber has fallen out of the specified range for both light and humidity.

### GMpi Software Dependencies

The scripts used by the GMpi for the two sensors, temperature and humidity (DHT22) and light (TSL2591), are dependent on the open source Python libraries *Adafruit_DHT* (github.com/adafruit/Adafruit_Python_DHT) and *python-tsl2591* (github.com/maxlklaxl/python-tsl2591). These libraries enable the devices to read and process output from the sensors. The camera uses software that is already packaged with the Raspbian Stretch operating system (I2C). To combine automation, remote monitoring, and data collection, we developed our own software package, *GMpi_Pack*, which is freely available on GitHub (https://github.com/BrandinGrindstaff/GMpi) under a GNU General Public License (v3). *GMpi_Pack* is composed of seven scripts: *GMPi_Pack.py* (the main library), *configuration.py, packageTest.py, picSnap.py, sense.py, upload.py*, and *install.sh*. After getting the software from GitHub, users should run the *install.sh* script. This script will attempt to download and install all software dependencies for *GMpi_Pack* automatically. Next, users should run the *configuration.py* script, which sets the minimum and maximum thresholds for light intensity and creates a configuration file allowing the end-user to easily modify important information needed for the GMpi to operate. The *packageTest.py* script is included for troubleshooting and gives the user a chance to test the device to confirm that the GMpi is running correctly. The three scripts, *sense.py, picSnap.py*, and *upload.py*, call functions from the main *GMPi_Pack.py* library to carry out sensor readings, capture images, and upload data to a cloud services, respectively. These scripts are intentionally independent to allow for modularity when scheduling “Cron jobs” to automate these processes. Cron jobs are setup using the Unix cron utility, a time-based job scheduler that comes packaged with Raspbian Stretch and other Unix-like operating systems. This allows the user to run shell commands on a time-based schedule, making it possible to run scripts automatically at user-set intervals.

### Protocol Feasibility

The protocol in the appendices provides the detailed instructions for setting up the GMpi and *GMpi_Pack*. They include networking, connecting the hardware, and installing the software. Networking will vary greatly between institutions and it is advisable to work with your IT department to work around firewalls that make remote monitoring and secure shell (SSH) access difficult. Both sensors utilized by the GMpi require soldering before use, however, there may be other options available at a slightly higher cost. For our use, the light sensor (TSL2591) required that the pin headers be soldered onto its printed circuit board, and the temperature and humidity sensor (DHT22) required a 10KO resistor be soldered between the power pin and data pin. The software we developed for the Raspberry Pi is dependent on open source tools that are available from Github and AdaFruit, as well as other online sources. Our software, and all of its dependencies, can be downloaded from GitHub (https://github.com/BrandinGrindstaff/GMpi) and set up using the scripts we have provided. To remotely alert users if the chamber or greenhouse falls below desired thresholds, the GMpi uses Webhooks to enable it to send alerts through the Slack application. As many groups are now communicating through Slack, we think this will provide a quick way for all members of a research program to be informed that a system is failing. We also provide an example script that will take output sensor data and plot it so users can visually assess the conditions of the environment they are monitoring (**Figure 4**). The camera on the GMpi can be coupled with machine learning algorithms, such as openCV, to allow for plant phenotyping similar to the Bellwether phenotyping platform developed by Fahlgren et al. (2015). Using the camera in combination with image uploading to cloud storage can also enable the user to diagnose disease or other issues with specimens in the growth chamber remotely from anywhere with internet access.

**Figure 4:**
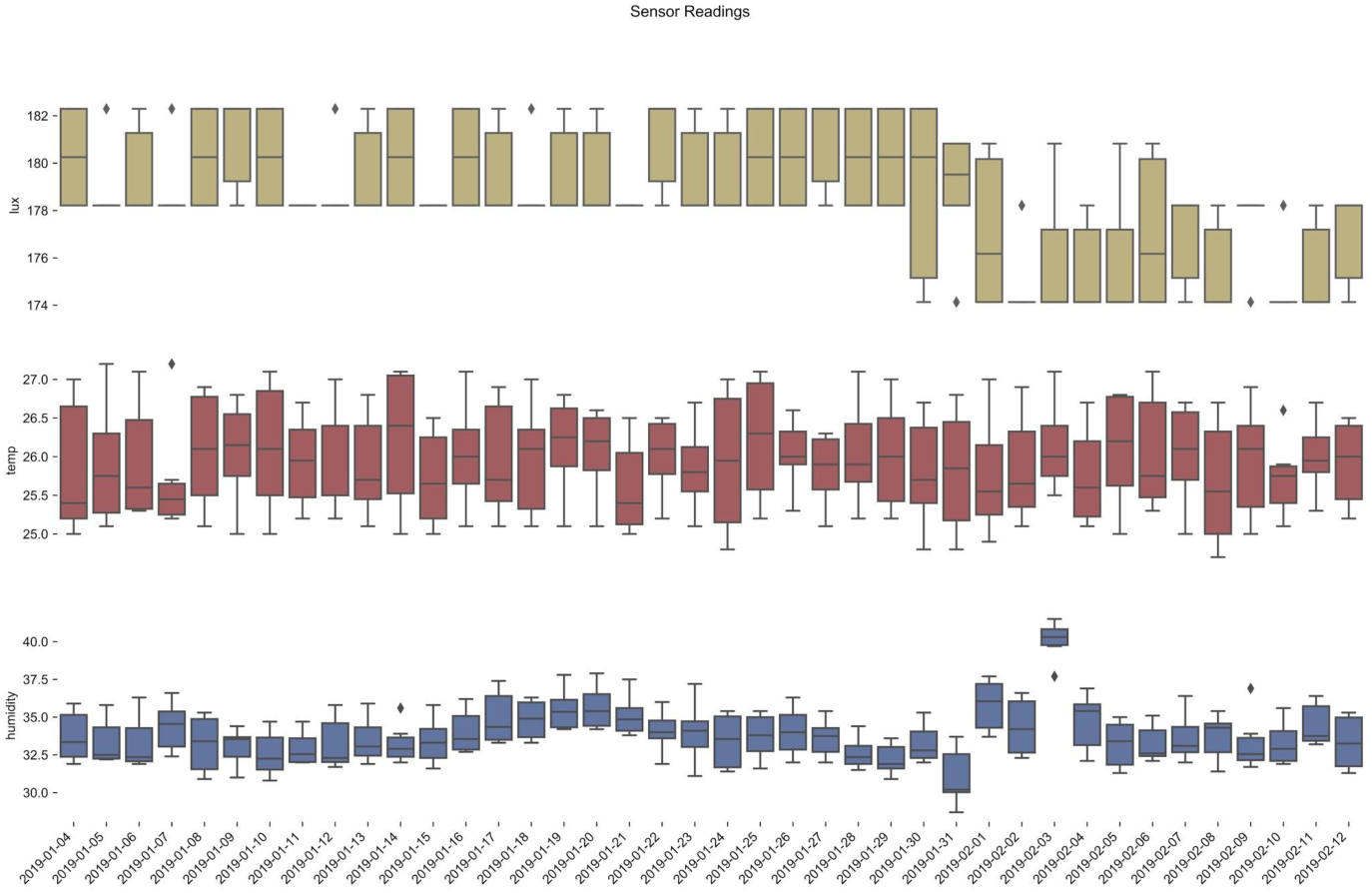
Plot of sensing data. Approximately one month of data captured with the GMpi. Graph displays light intensity (Yellow), temperature (Red), and humidity (blue) in box plot format.

## CONCLUSIONS

The GMpi gives researchers access to a low-cost option for environmental data monitoring and logging that can improve reproducibility of experiments and reliability of growth chamber conditions, as well as build large data sets that can be employed as phenotypic or environmental data in future studies. The Raspberry Pi computer also has a large developer community and is a great source of information for learning about its myriad applications. Due to its low cost and flexibility, the Raspberry Pi is becoming a popular tool for data collection and other tasks among researchers and industries. As its popularity as a tool grows, information and software available from the community should continue to improve in quality and volume. With a wealth of free, cost-effective, and open-source resources at hand, researchers are in an excellent position to leverage these tools to revolutionize plant science and improve reproducibility in experimentation with little impact on their budgets.

## ACKNOWLEDGMENTS

We would like to thank the Bond Life Science shop employees, Wayne Shoemaker, Danny Patterson, and Leon Toebben, for their help in soldering the sensors and Raspberry Pi computer together. We also thank Nick Valentine, Alan Marshall, and Jake Gotburg for their assistance and advice in working around and within the University of Missouri firewalls.

## APPENDIX 1.

### Raspberry Pi Hardware Setup

In Appendix 1 the physical setup of the device and its peripheries are covered, this includes the physical connection of the peripheries. With the exception of connecting the device to a monitor, the device should be powered off during every step of hardware setup.

#### Equipment Needed

Components ordered from www.adafruit.com

- Raspberry Pi Computer
  - Raspberry Pi Model 3B+ - 1.4GHz Cortex - A53 with 1GB RAM; Product ID: 3775
  - Adafruit Raspberry Pi B+ /Pi 2/ Pi 3 Case -Smoke Base; Product ID: 2258
  - 16GB SD/ MicroSD Memory Card −16GB Class 10; Product ID: 2693
  - 5V 2.4A Switching Power Supply with 20AWG MicroUSB Cable; Product ID: 1995
- Temperature and Humidity Sensor
  - DHT22 temperature - humidity sensor; Product ID: 385
- Light Sensor
  - Adafruit TSL2591 High Dynamic Range Digital Light Sensor; Product ID: 1980
- Camera
  - Raspberry Pi Camera Board v2 - 8 Megapixels; Product ID: 3099
  - Adafruit Raspberry Pi Camera Board Case with ¼” Tripod Mount; Product ID: 3253
  - Flex Cable for Raspberry Pi Camera −2 meters (If longer cable is desired); Product ID: 2144

Other important components that can be purchased elsewhere

- Female-female 2.54 to 2.0mm multi-colored jumper wires
- A computer with an SD card reader and internet access
- A monitor with an HDMI port for initial setup
- A keyboard and mouse that use a USB port to interface with the raspberry pi during initial setup
- An HDMI cable
- An antistatic wrist strap to protect your circuit boards from static discharge while handling
- Soldering Iron
- Heat shrink tubing or electrical tape

#### Connecting Sensors and Other Peripheries to the Raspberry Pi

The light intensity sensor (TSL2591) and the temperature and humidity sensor (DHT22) can be interfaced with the Raspberry Pi via GPIO (generalized purpose input/output) pins. They are the 40 pins that run along the upper part of the single board computer. The GPIO pins can deliver 3.3V or 5.0V of electricity, ground, and data transfer. The data transfer pins include both I2C (inter-integrated circuit) and UART (universal asynchronous receiver-transmitter) protocols, although the later is not used on the GMpi.

##### TSL2591 Light Intensity Sensor

1. Requires the included 6-pin header to be soldered to the device.
2. Connect female-female jumper wires to pins labeled on sensor “Vin”, “GND”, “SDA”, and “SCL” using a different color for each to distinguish between them easily when connected to the Raspberry pi. We suggest to use white or red for “Vin”, the power in connector, and black or brown for “GND”, the ground. This coloring scheme is standard convention, like the battery connectors of a motor vehicle.
  a. Connect the SDA (serial data) wire to GPIO pin 3 on the Raspberry Pi (**Figure 2**) and connect the SCL (serial clock) wire to GPIO pin 5. These two pins facilitate data transfer for this sensor which uses I2C serial protocol.
  b. Connect the ground wire to GPIO pin 9 and the power wire to GPIO pin 17, giving 3.3v power to the sensor and grounding it.

##### DHT22 Temperature Humidity Sensor

1. Connect jumper wires to the power, data, and ground pin, using a different color for each (same as above; adhering to the convention for power and ground if desired). Facing the grid upwards (front-up) the pins are from left to right power, data, an unused pin, and ground.
2. A 10kΩ resistor must be soldered between the power jumper wire and the data jumper wire in order to transmit data from the temperature and humidity sensor (DHT22) to the Raspberry Pi. Burn a small area of the wire casing off both jumper wires using a lighter. Solder the 10kΩ resistor between the power and data wire (**Figure 2**).
3. Cover the resistor and exposed wire with heat shrink wire sheathing. Heat the wire sheathing with a wire until it fits snugly. Alternatively it is also acceptable to use electrical tape so long as the circuit is insulated to prevent shorting.
4. Connect the power wire of the temperature and humidity sensor (DHT22) to GPIO pin 1, the ground wire to pin 6, and the data wire to pin 11 of the Raspberry Pi (**Figure 2**).

##### Camera and Camera Circuit Board

Connect the camera circuit board to the CSI-2 port in the lower middle of the Raspberry Pi (**Figure 2**). Remember to pull up the black tab on the port to insert the ribbon cable and to close the tab by pressing down the it. The camera listed above has a key to manually focus the camera. We suggest to wait to focus the lens of your camera till your GMpi is completely set up and your camera is in it’s final fixed position.

##### MicroSD Card

Insert the microSD card into slot on the bottom left of the Raspberry Pi. Before you do however, install the Raspbian Stretch operating system following the steps outlined in **Appendix 2**.

##### HDMI

Connect the HDMI cable between the Raspberry Pi and the HDMI compatible monitor for the initial setup, afterwards the GMpi can run “headless”, or without a monitor, using SSH (secure shell) protocol.

## APPENDIX 2.

### Raspberry Pi Initial Setup

In Appendix 2, we cover how to install the Raspbian stretch operating system onto the Raspberry Pi and configure it to use the GMpi software. This section also generally describes connecting the Raspberry Pi to a network, which will be different for every user depending on the network being used.

#### Raspberry Pi Operating System Installation

(version 4.14 of Raspbian Stretch was used, but the most recent version is recommended).

1. Download “Raspbian Stretch with Desktop” from “https://www.raspberrypi.org/downloads/raspbian/” *note: downloads as a .zip file.
2. Download “etcher” from “https://etcher.io”.
  a. select “Download for Windowsx64” (otherwise download the version compatible with your operating system).
3. Set up Etcher software.
  a. accept license agreement and install.
4. Run etcher software; select “year-month-day-raspian-stretch.zip” (that date is the date this version was released.
  a. Select the SD card you plan to use.
  b. Select the flash action and wait for the process to finish.
  c. Safely remove the SD card from your computer.
5. Plug in microSD card into your Raspberry Pi.
  a. Boot up (HDMI screen required for this step).

#### Configuring Raspberry Pi

(Instructions with a screen, not SSH)

1. Select raspberry symbol in the top left of the screen.
2. Scroll down to preferences.
3. Select “Raspberry Pi Configuration”.
4. Click the interfaces tab, and Enable…
  a. SSH, to connect wirelessly.
  b. I2C, for the light sensor.
  c. Camera to use the camera.
  d. Remote GPIO, to use our sensors over the internet.
5. Click Localization tab.
6. Select “Set WiFi Country; Scroll down and select your country.
7. Select “Set Time Zone” and select your timezone.
8. Select Keyboard, and select the corresponding locality/language that matches your keyboard configuration.
9. Click “OK”, and reboot the Raspberry Pi

#### Networking

In order to connect to the GMpi device remotely, you must use an HDMI compatible monitor, a mouse, and keyboard for the initial configuration to enable the SSH as outlined in the **Configuring Raspberry Pi** section of **Appendix 2**. With the device configured to use SSH, open the terminal, and enter the following command:

`ifconfig`

This command will display your network information on the terminal. In the ‘wlan0’ section, (the third block of text), look for the local IP address labeled ‘inet’. If your network has dynamic IP addresses, the address will change on a regular basis, requiring you to repeat this process with the new IP address. Therefore, setting up a static IP address is highly recommended. Universities also tend to have advanced network security protocols, for example, the University of Missouri uses WPA2 Enterprise wireless security protocol. Configuring the device may therefore also require advanced configuration changes to connect to the device wirelessly. Because of these reasons, it is recommended to contact your institution’s Information Technology department and request a static IP address, or other additional assistance with this step.

## APPENDIX 3

### Rclone and GMpi Software Installation Protocol

This section covers installation of the rclone software that gives the GMpi the ability to upload collected data to a cloud storage server as well as the installation of the GMpi software. It also covers the use of cron, a time based job scheduling software included with the Raspbian stretch operating system.

#### Installing and Configuring rclone for File Uploading

For more information access the rclone developer’s github (https://github.com/pageauc/rclone4pi). The installation protocol is based on the instructions provided by the developer.

1. Open the terminal and install rclone.
  a. `curl-L` https://raw.github.com/pageauc/rclone4pi/master/rclone-install.sh | `bash`
2. Upgrade to most recent version.
  a. `cd ∼/rpi-sync`
  b. `./rclone-install.sh upgrade`
3. Configure rclone.
  a. `rclone config`
  b. You will be given a list of options, select new remote by entering “n”.
  c. It will ask you to give an arbitrary name for your remote. It is important to remember this name because you will have to call it in the configuration file in order for it to work.
  d. You will then be given a list of file sharing services. Select the service of your choice. We decided to use the google drive service, but this is the end-users choice.
  e. <press enter> #this field is for locking program locally to a user name.
  f. <press enter> #this field is for setting a password to use this program locally.
  g. Select the type of access you want to give this program to the selected cloud account. Full access is recommended, select full access by entering “1”
  h. You will see a host of options, pressing enter without any other inputs will accept the default options.
    i. `y` #use autoconfig
  i. <press enter>
  j. The last input will make the program open your browser and take you to services login screen.
    i. Log in and select “allow” for rclone to run. It will ask you if you have a team drive, select “y” or “n”
  k. Finally it will give you an overview of your choices, look for errors now! If it looks right, type “`y` “
  l. Now you can Exit rclone config.
    i. `q`

#### GMpi Automatic Software Setup

Using the *install.sh* script, the end user is able to automatically download all libraries and software for the GMpi. If the end-user chooses to manually download the GMpi software and it’s dependencies follow the steps in the **GMpi Manual Software Setup** section, otherwise skip to the section titled **Scheduling Sensor Readings with Cron** after completing the steps below.

1. `git clone` https://github.com/BrandinGrindstaff/GMpi.git
2. `cd GMpi/`
3. `bash install.sh`
4. `python3 configure.py`

After running the *configure.py* script, a new file named ‘config.txt’ will be generated. Open this configuration file and replace all values listed as ‘<REPLACE>’ with the values you want to use for monitoring (e.g., times for when lights are on/off). The ranges specified here will also be used for alerting the user when growth chambers or greenhouses fall out of desired ranges for sensing.

#### GMpi Manual Software Setup

The *install.sh* script included with GMpi will install all software automatically, however if end-users wish to install software independently, steps are provided below. Use the following commands to download the GMpi software from github and create the configuration file ‘config.txt’.

1. `git clone` https://github.com/BrandinGrindstaff/GMpi.git
2. `cd GMpi/`
3. `python3 configure.py`

Follow the next two sets of instructions to manually install dependencies.

##### Humidity and Temperature Software Setup

For more information, go to https://github.com/adafruit/Adafruit_Python_DHT

1. Download/Install *Adafruit_Python_DHT* Library for the humidity and temp sensor.
  a. Open the terminal and download file.
    i. `git clone` https://github.com/adafruit/Adafruit_Python_DHT.git
  b. Move to the software folder
    i. `cd Adafruit_Python_DHT`
  c. Update and get dependencies in order for the software to work correctly.
    i. `sudo apt-get update`
    ii. `sudo apt-get upgrade`
2. Reboot the Raspberry Pi to allow the update to be recognized.
  a. `sudo reboot`
3. Install Python 3
  a. `sudo apt-get install build-essential python-dev python -open ssl`
  b. `sudo python3 setup.py install`

##### Light Sensor Software Setup

For more info on setup see https://github.com/maxlklaxl/python-tsl259 and https://learn.adafruit.com/adafruits-raspberry-pi-lesson-4-gpio-setup/configuring-i2c for configuring I2C.

1. Download/Install *python-tsl2951* Library for light sensor
  a. `git clone` https://github.com/maxlklaxl/python-tsl2591.git
2. Install dependency libffi to resolve compatibility issues with Python 3
  a. `sudo apt-get install libffi-dev`
3. Configure I2C (for light sensor)
  a. `sudo apt-get install -y python-smbus`
  b. `sudo apt-get install -y i2c-tools`
  c. `sudo reboot`
4. Wait for the Raspberry Pi to reboot, and then connect

#### Scheduling Sensor Readings with Cron

The cron utility is a job scheduling software included in most distributions of Unix-like operating systems that is included with the Raspbian Stretch operating system used with GMpi. It uses the time that is set on the Raspbian Stretch operating system (displayed in the bottom right of the screen, at the far right end of the task bar) to determine when to schedule a job, i.e. run a program. Cron can be extremely useful for automating process to run on a set time interval, the GMpi uses it to execute sensor reads, capture images, and upload data on set time intervals without manual execution by the end-user.

1. To create a Crontab file, in terminal execute the following commands
  a. `crontab -e`
    i. You will be given a set of options to select a text editor to make changes to the file, select nano, unless another option is more to your liking, and begin editing your crontab file.
    ii. A crontab accepts 6 fields separated by spaces.
      1. From left to right: minute, hour, day of month, month, day of week, and finally the command to be executed.
    iii. There is also a number of functions that can be done with integers in any of those first 5 fields so long as they are within the specified range of the field described in the text at the top of the file.
      1. To denote “first through last” in one of the first 5 fields use an asterisk.
    iv. Crontabs evokes the command from the user’s HOME directory, so it is important to include file pathing in your command.
      1. (1) For example
        a. `*/2 * * * * cd ∼/GMpi && python3 sense.py`
          i. This tells chrontabs to every other minute (`*/2`), of every hour (*) of every day of the week (*) of every month (*) and every day of the week (*), change directory (`cd`) from user’s HOME folder to GMpi (`∼/GMpi`) folder and (`&&`) use python3 to run “`sense.py`” and collect sensor readings.
      2. For more information read the text at the top of the created crontab file and/or read the manual to crontab or chron included with the Raspbian Stretch operating system.

### Remote Notifications

To create an incoming webhook for Slack notifications, first choose which workspace you wish to receive the alert. You may also wish to make the GMpi its own channel to communicate on. Once you have decided on or created a new workspace, you will need to install a new app. To do this click +add apps at the bottom of the left most column. Search for incoming webhooks and click install. Once installed you can then modify the settings to choose which channel you would like the GMpi to post to, add descriptive labels, and customize names and icons. The URL link included in the settings section is what you want to copy and replace in the configuration file.

